# Reduced somatostatin signalling leads to hypersecretion of glucagon in mice fed a high fat diet

**DOI:** 10.1101/2020.04.07.028258

**Authors:** Joely A. Kellard, Nils J. G. Rorsman, Thomas G. Hill, Sarah L Armour, Martijn van der Bunt, Patrik Rorsman, Jakob G. Knudsen, Linford J. B. Briant

## Abstract

Elevated plasma glucagon is an early symptom of diabetes, occurring in subjects with impaired glucose regulation. Here we explored alpha-cell function in female mice fed a high fat diet (HFD) – a widely used mouse model of pre-diabetes. *In vivo*, HFD-fed mice have increased fed plasma glucagon levels that are unaffected by elevation of plasma glucose. To explore the underlying mechanisms, we conducted experiments on isolated islets and in the perfused pancreas. In both experimental models, glucagon secretion under both hypo- and hyperglycaemic conditions was elevated. Because Ca^2+^ is an important intracellular regulator of glucagon release in alpha-cells, we fed mice expressing the Ca^2+^ indicator GCaMP3 specifically in alpha-cells the HFD. In mice fed a control (CTL) diet, increasing glucose reduced intracellular Ca^2+^ ([Ca^2+^]_i_) (oscillation frequency and amplitude). This effect was not observed in HFD mice where both the frequency and amplitude of the [Ca^2+^]_i_ oscillations were higher than in CTL alpha-cells. Given that alpha-cells are under strong paracrine control from neighbouring somatostatin-secreting delta-cells, we hypothesised that this elevation of alpha-cell output was due to a lack of somatostatin (SST) secretion. Indeed, SST secretion in isolated islets from HFD mice was reduced but exogenous SST also failed to suppress glucagon secretion and Ca^2+^ activity from HFD alpha-cells, in contrast to observations in CTL mice. These findings suggest that reduced delta-cell function, combined with intrinsic changes in alpha-cell sensitivity to somatostatin, accounts for the hyperglucagonaemia in mice fed a HFD.

## Introduction

Type 2 diabetes (T2D) is characterised by elevated circulating glucose. Lack of insulin plays an important role in the development of hyperglycaemia and glucose intolerance in T2D. However, it is also recognized that abnormal glucagon secretion contributes to the development of glucose intolerance and that T2D is best characterized as a bihormonal disorder [1, 2].

Glucagon is secreted from alpha-cells of the pancreatic islets when plasma glucose falls below ∼5 mM. Glucagon secretion is regulated within the islet by both intrinsic and paracrine mechanisms [3]. The mechanism(s) by which glucose inhibits glucagon secretion is still a matter of debate [3-10] but it is widely agreed that the effect is, at least in part, intrinsic to the alpha-cell. For example, glucose has been proposed to increase the intracellular ATP and that this, via closure of plasmalemmal ATP-regulated K^+^ (K_ATP_) channels, results in membrane depolarization and reduction in action potential height (due to voltage-dependent inactivation of the Na^+^ channels involved in action potential firing). This culminates in reduced activation of voltage-gated Ca^2+^ channels and, consequently, exocytosis of glucagon-containing granules [11]. However, glucagon release is also influenced by local paracrine signals. These include somatostatin [12, 13] and insulin [14, 15] from islet delta- and beta-cells, respectively.

In some patients with T2D, the normal relationship between plasma glucose and glucagon is reversed and hyperglycaemia stimulates rather than inhibits glucagon secretion [11, 16]. The dysregulation of glucagon secretion is detectable even prior to the onset of T2D; hyperglucagonaemia is observed in obese patients [17, 18] and patients with impaired fasting glycaemia [19]. Although it is clear that glucagon is central in the aetiology of T2D, we still do not understand how glucagon secretion is affected by the changes in whole body metabolism preceding the onset of the disease. In particular, the impact of high-fat diet (HFD) feeding – widely regarded as a model of pre-diabetes [20] – on glucagon secretion remains unknown. Exposure of whole islets to high levels of palmitate for up to 72h changes both insulin, glucagon and somatostatin secretion [21, 22] as well as whole-islet gene expression [23] and metabolism [24, 25]. Furthermore, isolated islets from HFD mice exhibit elevated glucagon secretion when exposed to high glucose concentrations [26]. However the mechanism by which this elevation occurs remains unresolved and obscured by the conflicting *in vivo* observations that circulating glucagon is increased [26], decreased [27] or unchanged [28] in HFD mice. Here we investigate the effects of HFD feeding on alpha-cell function and the paracrine regulation of glucagon secretion.

## Methods

### Ethics

Experiments were conducted in strict accordance to the UK Animals Scientific Procedures Act (1986) and University of Oxford ethical guidelines. All work was approved by the local Ethical Committee.

### Animals

Mice expressing GCaMP3 specifically in alpha-cells were generated by crossing *Gt(ROSA)26Sor*^*tm38(CAG-GCaMP3)Hze*^ mice (Jackson Laboratory No. 014538) with mice carrying an insert containing glucagon promoter-driven iCRE (*Tg(Gcg-icre)*^*12Fmgb*^ mice; see [29]). All mice were female and fully backcrossed to a C57BL/6J background. Unless otherwise indicated animals had *ad libitum* access to food and water. All animals were house in an SPF facility on a 12:12h light:dark cycle at 22°C. In all cases where animals were fasted, food was removed at 08.30 a.m. (30 min into the light phase). Immediately after weaning, mice were fed a high fat (HFD, % kcal: protein 18.3, carbohydrate 21.4, fat 60.3; TD.06414, Envigo) or a control diet (CTL, % kcal: protein 20.5, carbohydrate 69.1, fat 10.5; TD.08806 Envigo) for 12 weeks. Body weight was recorded every 14 days.

### Glucose tolerance test

Following 6 h of fasting, animals received an intraperitoneal (i.p.) injection of D-glucose (2 g/kg; IPGTT). Blood glucose concentrations were measured at 0, 15, 30, 60 and 120 min after the injection. A sample was also taken 15 mins prior to the injection (“Rest”). 25 µl of blood was obtained by tail vein puncture at 0 and 30 min in EDTA-coated capillary tubes. Whole blood was immediately mixed with 5 µl of aprotinin (1:5, 4 TIU/ml, Sigma-Aldrich UK) and kept on ice until it was centrifuged at 2600 g at 4°C. Plasma was then removed and stored at -80°C.

### Fed plasma measurements

Tail vein blood samples were also taken from *ad libitum* fed mice with free access to water, housed in their home cage. Blood samples were taken at 09:00, 13:00 and 17:00 and processed as described above.

### Insulin tolerance test

Following 4 h of fasting, animals received an i.p. injection of insulin dosed on total body weight (0.75 U/kg total body weight; Actrapid, Novo Nordisk). This insulin tolerance test (ITT) involved measuring blood glucose concentrations were measured at 0, 15, 30, 60 and 120 min after the injection. At fixed time points following the injection, 25 µl of blood was obtained and processed as above.

In an additional experiment, mice were given an insulin bolus where the insulin was dosed on lean mass. Initial experiments using EchoMRI™(EchoMRI LLC, USA) demonstrated that CTL mice were 69.5±2.1% lean mass, whereas HFD fed mice were 59.5±3.2% lean mass (p=0.023, n=6 CTL and 6 HFD mice, unpaired t-test; **Table 1**). Therefore, for the lean mass-based insulin injections, CTL mice received 0.75 U/kg total body weight, whereas HFD fed mice received 0.64 U/kg, thereby giving the mice the same dose of insulin per gram lean mass (1.08 U/kg lean mass).

**Table 1:** Body composition analysis of mice on HFD and CTL diet for 12 weeks by EchoMRI™

| Parameter | CTL (n=6) | HFD (n=6) | p |
| --- | --- | --- | --- |
| Weight (g) | 20.77±0.35 | 26.25±1.69 | 0.0035 |
| Fat % | 20.33±2.46 | 32.88±4.19 | 0.0273 |
| Lean mass % | 69.50±2.09 | 59.50±3.24 | 0.0268 |
| Total water % | 57.16±4.42 | 48.63±5.61 | 0.0151 |
| Fat (g) | 4.21±0.49 | 8.90±1.59 | 0.0183 |

### Islet isolation

Mice were culled by cervical dislocation. Pancreatic islets were isolated by liberase digestion followed by manual picking. Isolated islets were, pending the experiments, maintained in short-term (<24 h) tissue culture in RPMI 1640 (11879-020, Gibco, Thermo Fisher Scientific) containing 1% penicillin/streptomycin (1214-122, Gibco, Thermo Fisher Scientific), 10%FBS (F7524-500G, Sigma-Aldrich) and 11 mM glucose prior to the measurements.

### Static secretion experiments

Islets isolated from HFD and control mice were incubated in 11 mM glucose media. All secretion experiments were conducted on the day of islet isolation, following 1-2 h culture. Size-matched batches of 20 islets were then pre-incubated in 0.2 ml KRB (in mM; 140 NaCl, 5 KCl, 1.2 MgCl_2_, 2.6 CaCl_2_, 1 NaH_2_PO_4_, 5 NaHCO_3_, 10 HEPES (pH 7.4)) with 2 mg/ml BSA (S6003, Sigma-Aldrich) and 3 mM glucose for 1 hour at 37 °C. Following this, islets were subjected to 1 mM or 6 mM glucose KRB with 0.2% BSA for 1 hour. The supernatant was removed, quickly frozen and stored at -80 °C. For measurement of total glucagon and insulin contents, the islets were lysed in HCl:ethanol (1:15) at the end of the experiment, sonicated and stored at -80 °C.

### The *in situ* perfused mouse pancreas

Briefly, the aorta was ligated above the coeliac artery and below the superior mesenteric artery and then cannulated. The pancreas was perfused with KRB containing varying concentrations of glucose and somatostatin as indicated in the figures, at a speed of 1.34 µl/min/mg pancreas weight using an Ismatec Reglo Digital MS2/12 peristaltic pump. Pancreatic weight was estimated from whole body weight as previously described [30, 31]. The perfusate was maintained at 37°C using a Warner Instruments temperature control unit TC-32 4B in conjunction with an in-line heater (Warner Instruments P/N 64-0102) and a Harvard Apparatus heated rodent operating table. The effluent was collected in intervals of 1 min into 96-well plates kept on ice and containing aprotinin. Samples were subsequently stored at -80°C pending analysis of glucagon content.

### Hormone measurements

Plasma insulin and glucagon was determined using insulin and glucagon mouse sandwich ELISA (10-1113-01 and 10-1281-01 from Mercodia, Sweden). Insulin and glucagon concentrations from *ex vivo* islet experiments were measured using mouse/rat insulin-glucagon sandwich ELISA (K15145C, Mesoscale Discovery, USA) and somatostatin concentration was determined using radioimmunoassay (Life Science AB, Sweden). Glucagon concentrations from the perfusate of the *in situ* perfused mouse pancreas were measured using the U-plex Glucagon ELISA (K1515YK, Mesoscale Discovery). All measurements were conducted according to the manufacturers’ protocols.

### GCaMP3 imaging and calculation of [Ca^2+^]_i_ spike frequency and amplitude

Time-lapse imaging of the intracellular GCaMP3 was performed on the inverted Zeiss AxioVert 200 microscope, equipped with the Zeiss LSM 510-META laser confocal scanning system, using a 40x/1.3 NA objective. The chamber was continuously perfused at a rate of 200 μl/min with KRB solution (described above) containing 2 mg/ml BSA (S6003, Sigma-Aldrich), glucose and other compounds as indicated in the figures. All solutions were corrected for osmotic differences with mannitol [32]. GCaMP3 was excited at 488 nm and fluorescence emission collected at 530 nm at a frequency of 1.28Hz. Fiji (http://fiji.sc/Fiji) was used to identify and measure the intensity of the GCaMP3 signal in individual regions of interest (cells) over time. Spikes in GCaMP3 were manually annotated using Spike2 (http://ced.co.uk/) and defined as having a peak that was >20% of the signal:noise. Spike frequency was calculated from this annotated data, and spike amplitude was calculated from the spike-triggered average of the fluorescence signal over a fixed time window (10 time-steps).

### Perforated patch-clamp recordings

Islets isolated from chow-fed as well as CTL and HFD mice were used for electrophysiological recordings. These recordings (in intact islets) were performed at 33-34 °C using an EPC-10 patch-clamp amplifier and PatchMaster software (HEKA Electronics, Lambrecht/Pfalz, Germany). Unless otherwise stated, recordings were made in 3 mM glucose, to mimic hypoglycaemic conditions in mice. Currents were filtered at 2.9 kHz and digitized at > 10 kHz. A new islet was used for each recording. Membrane potential (V_m_) recordings were conducted using the perforated patch-clamp whole-cell technique as previously described [33]. The pipette solution contained (in mM) 76 K_2_SO_4_, 10 NaCl, 10 KCl, 1 MgCl_2_·6H_2_0 and 5 Hepes (pH 7.35 with KOH). For these experiments, the bath solution contained (mM) 140 NaCl, 3.6 KCl, 10 Hepes, 0.5 MgCl_2_·6H20, 0.5 Na_2_H_2_PO_4,_ 5 NaHCO_3_ and 1.5 CaCl_2_ (pH 7.4 with NaOH). Amphotericin B (final concentration of 25 mg/mL, Sigma-Aldrich) was added to the pipette solution to give electrical access to the cells (series resistance of <100 MΩ). Alpha-cells were confirmed by the presence of GCaMP3 or RFP. In WT islets, alpha-cells were identified by the presence of action potential activity at 3 mM glucose and ion channel properties [34]. In some recordings, GCaMP3 was also simultaneously recorded with a Hamamatsu ORCA 2, operated with MicroManipulator.

The frequency of action potential firing was calculated in MATLAB v. 6.1 (2000; The MathWorks, Natick, MA). In brief, a peak-find algorithm was used to detect action potentials. This was then used to calculate firing frequency and correlate average firing frequency (calculated every 1 s) with the GCaMP3 signal. Spike-triggered average waveforms were determined in Spike2 (CED, Cambridge, UK).

### Immunofluorescent staining

Whole pancreases were harvested and fixed in 4% PFA for up to 24 hours before embedding in wax, 5 µm thick sections were cut and stained using the antibodies indicated below.

#### Primary and Secondary antibodies used for staining of mouse pancreases. All dilutions are 1:500

| Ig (primary) | Ig (secondary) |
| --- | --- |
| Chicken anti GFP (Life Technologies, A10262) | Goat anti chicken 488 nm (Invitrogen, A11029) |
| Mouse anti glucagon (Sigma-Aldrich, G2654) | Goat anti mouse 568 nm (Invitrogen, 11032) |
| Guinea pig anti insulin | Goat anti guinea pig 633 nm |
| Rabbit anti somatostatin | Goat anti rabbit 488 nm |
| Rabbit anti SSTR2 (Abcam, ab134152) | Vector Labs DyLight Kit (DK-1488) |

Stained sections were imaged using a BioRad Radiance 2000 Scanning Laser Confocal Microscope.

### Statistics

All data are reported as mean±SEM, unless otherwise stated. Statistical significance was defined as p<0.05. All statistical tests were conducted in Prism 8.0 (GraphPad Software, San Diego, CA). For two groupings, a *t*-test was conducted. If the data were non-parametric, a Mann-Whitney test was conducted. For more than two groupings, a one-way ANOVA was conducted. If there were two independent variables, a two-way ANOVA was conducted. If the data passed normality criteria (D’Agostino’s test of normality and Bartlett’s test of equal variances), a parametric test was conducted with the appropriate post hoc test (Tukey or Sidak). If the normality criteria were not met, a Kruskal–Wallis test with Dunn’s multiple comparison test was conducted.

## Results

### HFD alters glucose homeostasis and plasma glucagon concentration *in vivo*

Following weaning, female mice were fed either a high-fat diet (HFD; 60% dietary calories from fat) or a control diet (CTL; 10% calories from fat) for 12 weeks. HFD feeding resulted in an increase body weight and fat mass (**Fig. 1a-b** and **Table 1**). To determine whether HFD feeding affected glycaemia, we measured blood glucose and plasma glucagon in *ad libitum* fed animals over several time points during the light phase. In HFD mice, blood glucose was not different between CTL and HFD mice at any time (**Fig. 1c**). Despite this, HFD mice had higher plasma glucagon levels than CTL mice in the beginning of the light phase (09:00 am; **Fig. 1d**). We analysed the glucagon:glucose ratio from all time points and found that it was higher in HFD mice (**Fig. 1e**), supporting the notion that the relationship between glucagon and glucose was altered in HFD mice. Insulin is a known paracrine inhibitor of glucagon [2, 14, 15], but levels of circulating insulin were in fact elevated in response to HFD feeding (**Fig. 1f**), making it unlikely that the elevated plasma glucagon in HFD mice was secondary to reduced plasma insulin.

**Figure 1:**
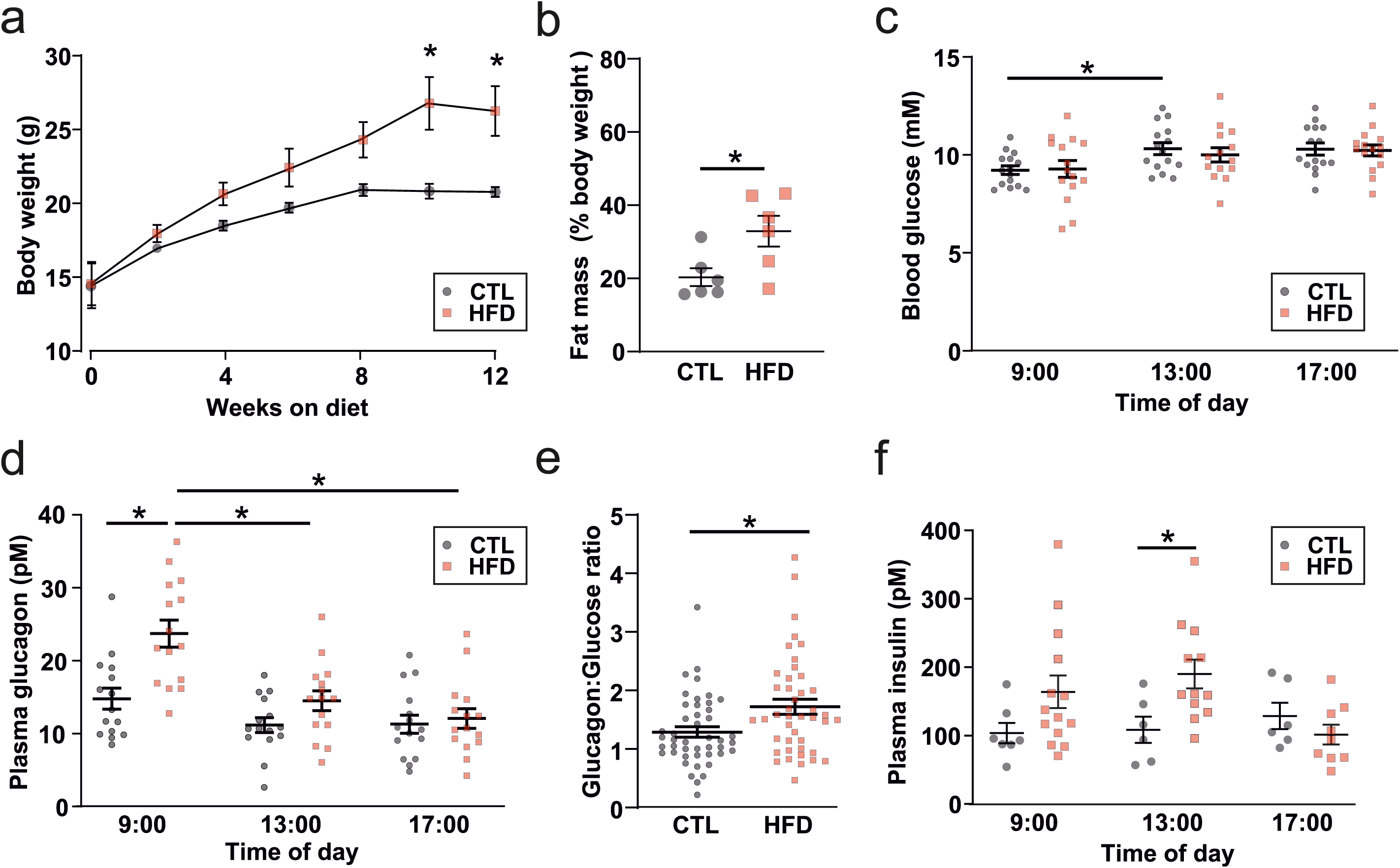
HFD feeding evokes hyperglucagonaemia *in vivo*. *a*. Increase in bodyweight in response to high fat diet (HFD) or control diet (CTL). Two-way repeated measures ANOVA, p<0.05=*. Data is presented as mean±SEM. *b*. Total body fat as % of total body weight in response to CTL or HFD at 12 weeks on diet. Unpaired t-test, p<0.05=*. Data is presented as mean±SEM. *c*. Fed blood glucose during the day for mice on the CTL (N=15) or HFD (N=14). Two-way repeated measures ANOVA, p<0.05=*. Data is presented as mean±SEM. *d*. Same as *c*. but plasma glucagon. Two-way repeated measures ANOVA, p<0.05=*. *e*. The Glucagon:Glucose ratio for mice in the CTL and HFD. Data taken for all 3 time points from *c*. and *d*. Unpaired t-test, p<0.05=*. Data is presented as mean±SEM. *f*. Same as *c*. but plasma insulin, CTL (n=6-7 mice), HFD (n=9-14 mice). Two-way repeated measures ANOVA, p<0.05=*. Data is presented as mean±SEM.

An increase in plasma glucose during a glucose tolerance test reduces circulating glucagon. This suppression is impaired in diabetic patients [35, 36] and conditions of impaired fasting glycaemia [37, 38]. HFD mice had impaired glucose tolerance and plasma glucose concentration was increased from 15 mM to 25 mM at 30 min (**Fig. 2a**). Plasma glucagon was reduced to the same extent as in CTL mice (**Fig. 2b**).

**Figure 2:**
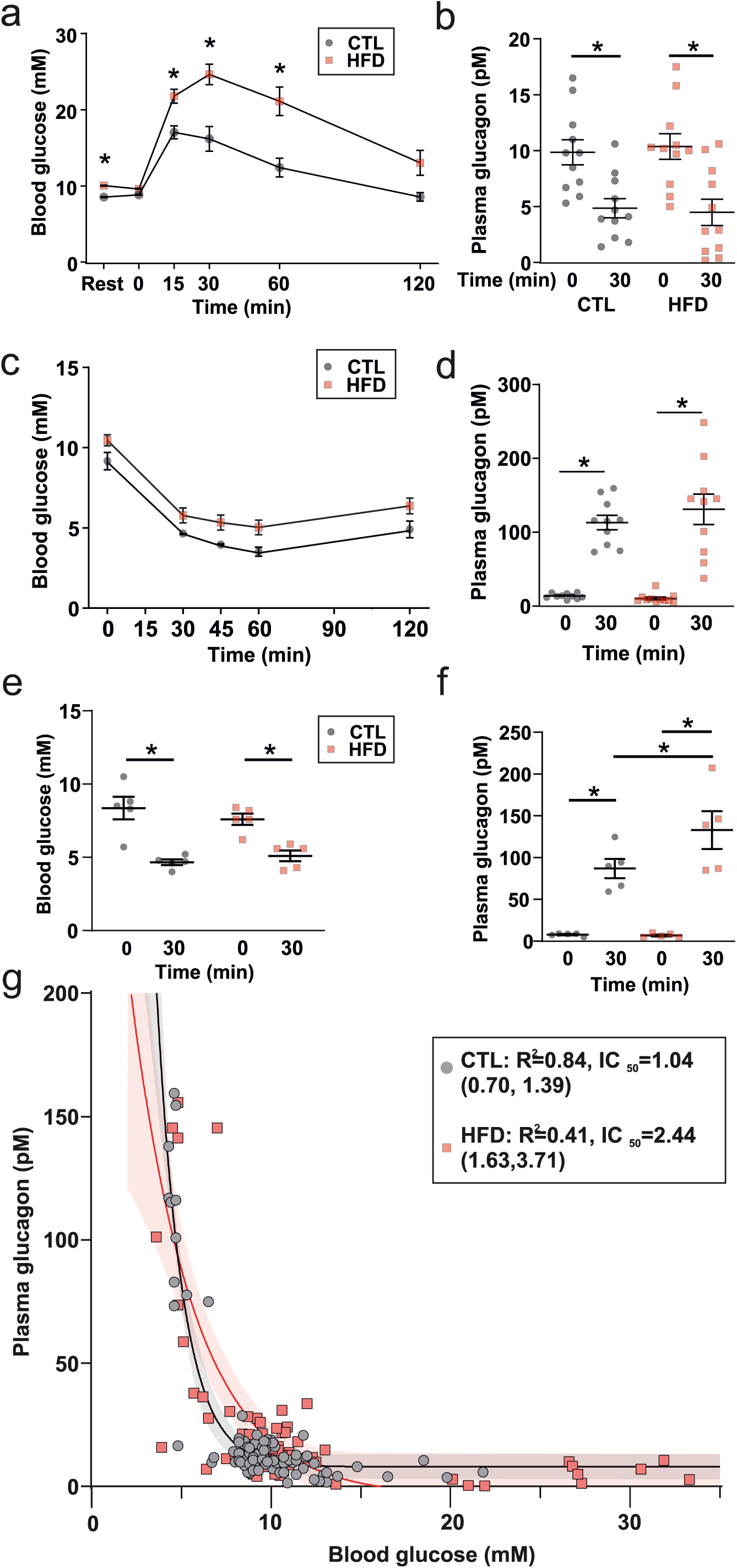
Counter-regulatory glucagon secretion is elevated in response to HFD feeding. *a*. Blood glucose of mice in response to an i.p. glucose tolerance test (GTT; 2 mg/kg) in mice on a CTL (n=12) or HFD (n=13) diet. Two-way repeated measures ANOVA, p<0.05=*. Resting value (Rest) is following a 6 h daytime fast. Data is presented as mean±SEM. *b*. Glucagon data from *a*. Two-way repeated measures ANOVA, p<0.05=*. n=11 for each diet. Data is presented as mean±SEM. *c*. Blood glucose following an i.p. insulin tolerance test (ITT) in mice fed a control (CTL, n=5) or high fat diet (HFD, n=5). Insulin was dosed based on total body weight (0.75 U/kg). Data is presented as mean±SEM. *d*. Plasma glucagon for data in *a*. for n=10 CTL and n=11 HFD mice. Two-way repeated measures ANOVA, p<0.05=*. Data is presented as mean±SEM. *e*. Blood glucose following an ITT dosed on lean mass (1.08 U/kg lean mass) n=5 CTL and n=5 HFD mice. Two-way repeated measures ANOVA, p<0.05=*. Data is presented as mean±SEM. *f*. Plasma glucagon for data in *c*. Two-way repeated measures ANOVA, p<0.05=*. *g*. Plot of blood glucose *vs* plasma glucagon for all *in vivo* data (including at rest, during GTT and during i.p. ITT dosed on total body weight). A single phase decay exponential (*A exp (- a [G])*; parameters *A* and *a*; *[G]* is the plasma glucose concentration) was fit to the CTL data (R^2^= 0.83) and HFD data (R^2^=0.41). The ‘half-life’ (t_1/2_) of the exponential decay is the glucose required to half glucagon secretion, and is therefore analogous to IC_50_. For CTL this was 1.04 (95% CI: [0.70, 1.39]) mM glucose and for HFD this was 2.44 [1.63, 3.71] mM glucose (N=91 CTL and N=90 HFD mice).

As glucagon is a counter-regulatory hormone, we also explored whether glucagon was inappropriately secreted in HFD fed mice during insulin-induced hypoglycaemia during an insulin tolerance test (ITT; **Fig. 2c**). When insulin was dosed according to total body weight, there was no difference in absolute glucagon (**Fig. 2d**).

During an ITT, the majority of glucose is taken up by skeletal muscle [39]. In the ITT (**Fig. 2c**) the insulin dose was calculated based on total body weight. However, because of the drastic difference in body composition between the diets (**Table 1**), this leads to an artificially high insulin dose in HFD mice. To understand whether the higher insulin bolus in the HFD mice resulted in a greater suppression of glucagon during the ITT, we also dosed insulin based on estimated lean body mass (**Fig. 2e-f**). Blood glucose levels were similar in the two groups 30 min after the insulin bolus, but plasma glucagon was elevated more in response to insulin in HFD than CTL mice (**Fig. 2e-f**), suggesting that glucagon is inappropriately increased in response to hypoglycaemia in HFD fed mice.

Finally, to understand how the relationship between glucose and glucagon was changed with HFD feeding, we combined all glucose and glucagon data from the ITT (dosed on total body weight) and GTT experiments (**Fig. 2g**). This demonstrated that *in vivo* glucagon closely follows an exponential relationship with glucose in CTL mice (R^2^=0.84, IC_50_ = 1.04 mM glucose). The relationship was markedly different in HFD fed mice (R^2^=0.40, IC_50_ = 2.44 mM) with a greater-than doubling in the glucose concentration required to suppress plasma glucagon by 50%. These data also clearly demonstrate that glucagon is inadequately suppressed by hyperglycaemic conditions (> 7 mM) – a known defect of TDM [8, 40].

### Intrinsic effects in islets drive the elevated plasma glucagon in HFD fed mice

To determine whether the elevated plasma glucagon was due to changes intrinsic to the islet, we measured glucagon secretion from isolated islets as well as from the *in situ* perfused mouse pancreas. In the perfused pancreas, glucagon secretion evoked by lowering plasma glucose from 6 to 1 mM was higher in HFD than in CLT mice (**Fig. 3a-b**). The increased glucagon secretion was also observed in static incubations of isolated islets exposed to 1 and 6 mM glucose (**Fig. 3e-f**). Finally, insulin secretion from the perfused mouse pancreas (**Fig. 3c-d**) and isolated islets (**Fig. 3g-h**) was (if anything) slightly (but non-significantly) elevated in HFD fed animals, reinforcing the conclusion from the *in vivo* experiments that the hypersecretion of glucagon was not due to reduced paracrine signaling from neighbouring beta-cells.

**Figure 3:**
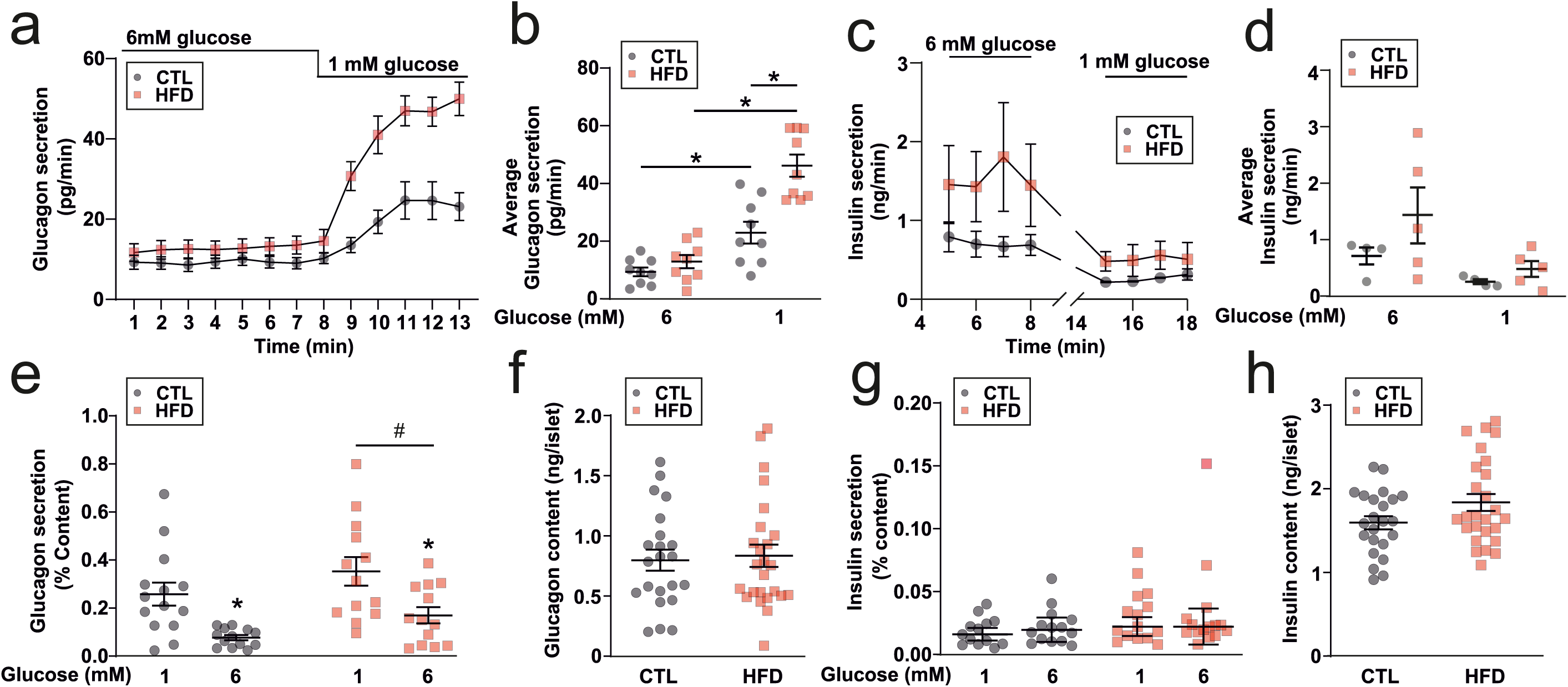
Glucagon secretion from *ex vivo* islets or the *in situ* perfused mouse pancreas is elevated in response to HFD. *a*. Glucagon measured in the perfusate of the perfused mouse pancreas from mice fed a control (CTL) or high fat diet (HFD). n=8-9 mice in each group. Two-way repeated measures ANOVA, p<0.05=*. Data is presented as mean±SEM. *b*. Data from *a*. but average steady-state values over each condition. Two-way repeated measures ANOVA, p<0.05=*. Data is presented as mean±SEM. *c*. Steady-state insulin measured in the perfusate of the perfused mouse pancreas from mice fed a control (CTL) or high fat diet (HFD). n=4 CTL and n=5 HFD mice. Two-way repeated measures ANOVA (significant source of variation: diet, p=0.21; time, p=0.012; interaction, p=0.22). Data is presented as mean±SEM. *d*. Data from c. but average steady-state values over each condition. Two-way repeated measures ANOVA, p>0.2. Data is presented as mean±SEM. *e*. Glucagon secreted from isolated islets from CTL and HFD mice (n=13 replicates from 6 mice). Two-way repeated measures ANOVA. Although there was no difference within a glucose concentration according to post-hoc analysis, there was an overall main effect between the diets (p<0.05=*). Data is presented as mean±SEM. *f*. Glucagon content from isolated islets from CTL and HFD mice (n=25 replicates from 6 mice). Unpaired t-test (p=0.92). Data is presented as mean±SEM. *g*. Same as c. but insulin secretion (n=15 replicates from 6 mice). Two-way repeated measures ANOVA, p<0.05=*. Data is presented as mean±SEM. *h*. Same as *d*. but insulin content (n=24 replicates from 6 mice). Unpaired t-test (p=0.51). Data is presented as mean±SEM.

### Alpha-cell dysfunction is associated with altered intracellular Ca^2+^ signaling

Electrical activity is an important determinant of alpha-cell glucagon output. We first conducted patch-clamp electrophysiology of alpha-cells from CTL and HFD mice and observed that HFD alpha-cells exhibit increase action potential firing (**Fig. 4a-e**). However, it was clear from these recordings that alpha-cells are heterogeneous in their electrical activity, with some alpha-cells displaying periods of quiescence in response to high (6 mM) glucose, and others responding with more subtle changes in frequency and/or action potential amplitude. Furthermore, the perforated patch-clamp technique is low throughput and restricts the study of alpha-cells to those on the outer layer of the islet. Alpha-cells exhibit oscillations in intracellular Ca^2+^ ([Ca^2+^]_i_) and changes in [Ca^2+^]_i_ drive glucagon secretion [41]. We performed parallel measurements of electrical activity and [Ca^2+^]_i_ in islets from mice expressing the genetically encoded Ca^2+^ indicator GCaMP3 under the *Gcg* promoter. We confirmed that electrical activity is correlated with [Ca^2+^]_i_ activity (**Fig 4f-h**). We conducted a cross-correlation of instantaneous firing frequency (calculated over a 2 second window) with the GCaMP3 signal, and observed them to be highly correlated (R^2^=0.6, **Fig. 4g-h**), demonstrating that [Ca^2+^]_i_ serves as a high-throughput proxy for electrical activity. We then ascertained that GCaMP3 was correctly targeted to the alpha-cells; we found that 84 ± 2% (n=3 mice) *GCG*^*+*^ cells expressed GCaMP3 (**Fig. 5a**). We then fed mice expressing GCaMP3 in alpha-cells a CTL or HFD. In islets isolated from CTL mice, spontaneous [Ca^2+^]_i_ oscillations were observed at 1 mM glucose that were suppressed in frequency and amplitude when glucose was increased to 6 mM glucose (**Fig. 5b-c and e**). However, there was no ‘typical’ alpha-cell Ca^2+^ signature; the changes in both frequency and amplitude were extremely variable. In HFD islets, [Ca^2+^]_i_ oscillations were also observed at 1 mM glucose but these were much less affected by elevation of glucose to 6 mM (**Fig. 5b-c**). In both CTL and HFD islets, the frequency of [Ca^2+^]_i_ oscillations was reduced by increasing glucose from 1 to 6 mM (**Fig. 5c**). However, the median frequency in 6 mM glucose was (on average) 2 fold higher in HFD than CTL alpha-cells. Furthermore, a larger proportion of alpha-cells remained active at 6 mM glucose in islets from HFD fed compared to CTL fed mice (**Fig. 5d**). It is notable than in CTL islets, ∼60% of alpha-cells remained active at 6 mM glucose (albeit at an extremely low oscillation frequency). Although glucose suppressed [Ca^2+^]_i_ oscillation amplitude in both CTL and HFD alpha-cells, alpha-cells from HFD fed mice had higher spike amplitudes than CTL mice at both 1 and 6 mM glucose (**Fig. 5e**).

**Figure 4:**
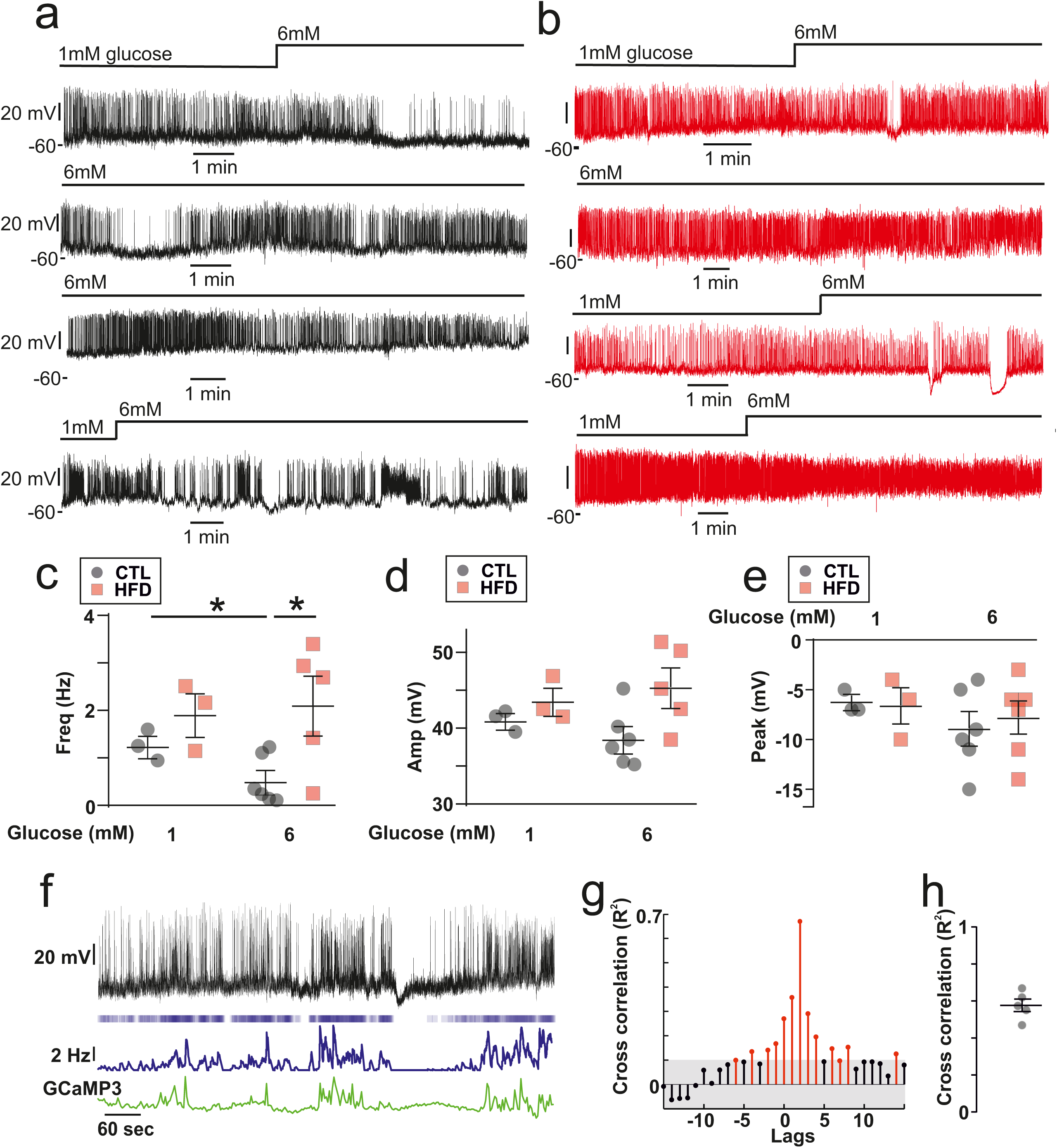
Electrical activity in CTL and HFD alpha-cells. *a*. Perforated patch-clamp recording of membrane potential (V_m_) from 4 alpha-cells from CTL mice. Note the variability in the response to high (6 mM) glucose. 6G = 6 mM glucose, 1G = 1 mM glucose. *b*. Recording of V_m_ from a HFD 4 alpha-cells from HFD mice. Note the variability in the response to high (6 mM) glucose. *c*. Firing frequency in 1 and 6 mM glucose (n=3-6 cells from N=3 CTL mice, N=3 HFD mice). Unpaired t-test, p<0.05=*. *d*. Action potential amplitude in 1 and 6 mM glucose (n=3-6 cells from N=3 CTL and N=3 HFD mice). Unpaired t-test, all p>0.11. *e*. Peak potential in 1 and 6 mM glucose (3-6 cells from N=3 CTL and N=3 HFD mice). Unpaired t- test, all p>0.3. *f*. Dual recording of GCaMP3 and V_m_ from an alpha-cell from a standard rodent chow-fed mouse. The upper trace is V_m_, below is a raster plot of action potentials with average firing frequency (calculated over a 2 s interval) and then the GCaMP3 signal from this cell. Note the correlation in average firing frequency and GCaMP3. *g*. Cross correlation of GCaMP3 signal and average firing frequency. The threshold for the cross correlation being deemed significant was R^2^=0.1. The lags was 1 time step (2 s). *h*. Maximum cross correlation from 5 alpha-cells from N=2 mice.

**Figure 5:**
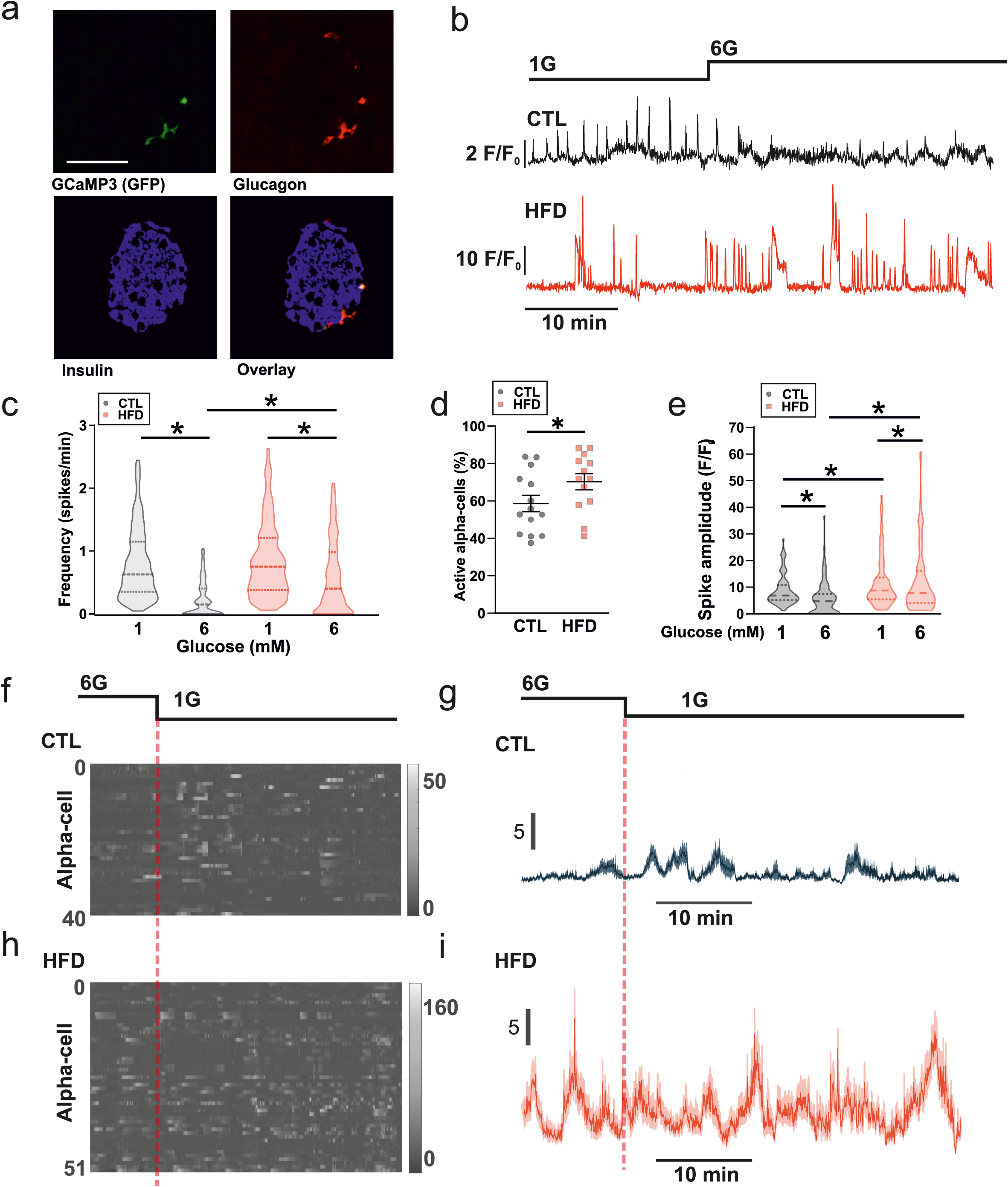
Alpha-cells from HFD mice exhibit elevated [Ca^2+^]_i_ oscillations *ex vivo*. *a*. Staining of GCaMP3, glucagon, insulin and an overlay in islets from *Gcg*^Cre-/+^ x floxed GCaMP3 mice. Representative of 65 cells from 3 mice. The % of cells that were GCaMP3+ and Gcg+ was 84±2%. Data is presented as mean±SEM. *b*. Glucose-dependent intracellular Ca^2+^ (GCaMP3; [Ca^2+^]_i_) signals in an alpha-cell from a mouse fed a control diet (CTL) and high fat diet (HFD). 1G = 1 mM glucose, 6G = 6 mM glucose. *c*. Frequency of [Ca^2+^]_i_ oscillations in response to 1 and 6 mM glucose. (n=508 cells from 7 CTL mice, n=561 cells from 7 HFD mice). Data are shown as median±quartiles. Two-way repeated measures ANOVA, p<0.05=*. Significance for HFD and CTL at 1 mM glucose is p=0.092. *d*. % of alpha-cells exhibiting [Ca^2+^]_i_ oscillations in CTL and HFD islets at 6 mM glucose. Unpaired t- test, p<0.05=*. Data is presented as mean±SEM. *e*. [Ca^2+^]_i_ spike amplitude in response to 1 and 6 mM glucose. (n=125 cells from 4 CTL and HFD mice). Data are shown as median±quartiles. Two-way repeated measures ANOVA, p<0.05=*. *f*. Raster plot of [Ca^2+^]_i_ signal from 40 alpha-cells from an islet from one CTL mouse. *g*. Average (±SEM) [Ca^2+^]_i_ response for all alpha-cells shown in *f*. *h*. Raster plot of [Ca^2+^]_i_ signal from 51 alpha-cells from an islet from one HFD mouse. *i*. Average (±SEM) [Ca^2+^]_i_ response for all alpha-cells shown in *i*.

Finally, in CTL islets we also observed the emergence of a degree of synchronicity of alpha-cell [Ca^2+^]_i_ in response to glucose (**Fig. 5f-I**; see also **Supplementary Fig. 1**) – an observation which has previously been attributed to somatostatin secretion from islet delta-cells [42] which may in turn be driven by beta-cells [14, 43]. In HFD islets, this synchronicity was unchanged by glucose.

### HFD islets exhibit somatostatin resistance and impaired somatostatin secretion

Alpha-cells are under strong paracrine regulation from neighbouring somatostatin-secreting delta- cells [43, 44]. Long-term exposure *in vitro* of islets to the non-esterified fatty acids oleate or palmitate has been shown to reduce somatostatin (SST) secretion [22]. We therefore hypothesized that the increase in glucagon secretion and [Ca^2+^]_i_ oscillatory activity at 6 mM glucose may be due to lowered somatostatin (SST) secretion. Indeed, glucose-stimulated SST secretion was 30% lower in islets isolated from HFD fed animals at both 1 mM and 15 mM glucose (**Fig. 6a**).

**Figure 6:**
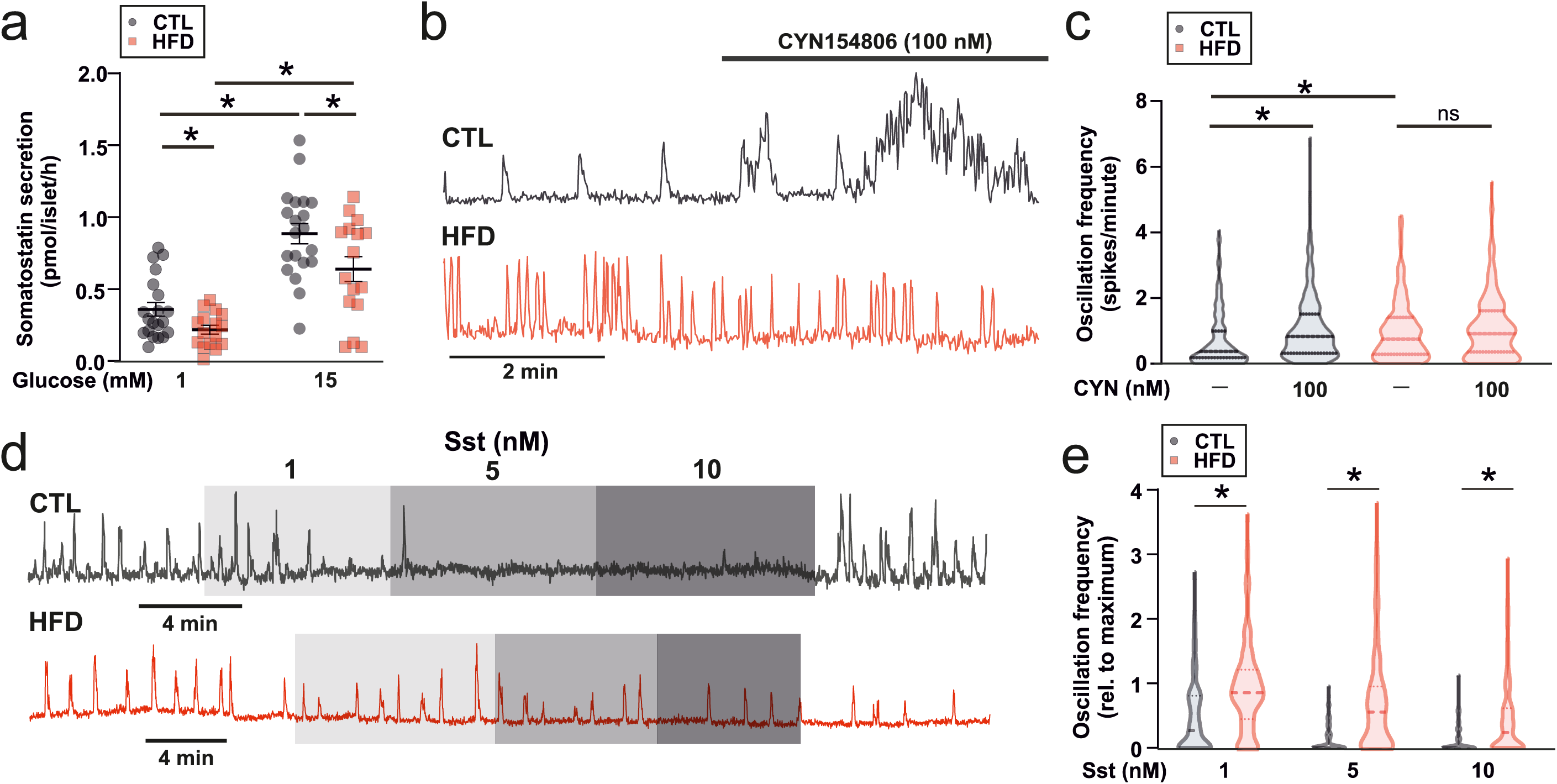
HFD results in changes in SST secretion. *a*. Somatostatin (Sst) secretion from islets isolated from control (CTL) and high fat diet (HFD) fed mice (n=16 replicates from 5 HFD and 5 CTL mice). T-test; p<0.05=*. Data is presented as mean±SEM. *b*. [Ca^2+^]_i_ signal (GCaMP3) from an alpha-cell from a CTL and HFD mouse in response to the SSTR2 antagonist CYN154806 (100 nM). Recording in 6 mM glucose. *c*. Average alpha-cell [Ca^2+^]i oscillation frequency in CTL and HFD islets in response to the SSTR2 antagonist CYN154806 (100 nM). Recording in 6 mM glucose. n=224 cells from 5 CTL mice and n=220 cells from 5 HFD mice. Data are shown as median±quartiles. *d*. [Ca^2+^]_i_ signal (GCaMP3) from an alpha-cell from a CTL and HFD mouse in response to 1, 5 and 10 nM Sst. Recording in 1 mM glucose. *e*. Average alpha-cell [Ca^2+^]_i_ oscillation frequency in CTL and HFD islets in response to 1, 5 and 10 nM SST. n=192 cells from 3 CTL mice and n=233 cells from 5 HFD mice. Two-way RM ANOVA; p<0.05=*. Data are shown as median±quartiles.

To determine whether the reduced SST secretion explained the lack of inhibition of glucagon secretion by glucose, we compared [Ca^2+^]_i_ oscillatory activity in CTL and HFD mice before and after pharmacological inhibition of somatostatin signalling in the islet. Mouse alpha–cells express primarily SST receptor 2 (SSTR2; see [7] and [45]). We therefore inhibited SST signaling using the SSTR2 inhibitor CYN154806 (CYN) and measured [Ca^2+^]_i_ oscillation frequency. At 6 mM glucose (a concentration associated with stimulation of somatostatin secretion in mouse islets; see Walker *et al*. [46]), addition of CYN increased [Ca^2+^]_i_ oscillation frequency significantly in alpha-cells from CTL but not in HFD mice (**Fig. 6b-c**). We also tested the capacity of exogenous SST to suppress alpha-cell [Ca^2+^]_i_ oscillation at 1 mM glucose. Whereas SST had a strong inhibitory effect in CTL islets, the effect was much weaker in HFD islets (**Fig. 6d-e**). In CTL islets, SST produced a concentration-dependent suppression of [Ca^2+^]_i_ oscillatory activity but this effect was less pronounced in HFD islets where [Ca^2+^]_i_ oscillations persisted at the high SST concentration tested (10 nM).

Collectively the effects of CYN154806 and exogenous somatostatin on [Ca^2+^]_i_ indicate that the alpha cells have become resistant to SST. We further explored this possibility using the perfused mouse pancreas. We first determined the IC_50_ of SST-induced suppression of glucagon to be 21 pM in chow- fed WT mice (**Fig. 7a**). Accordngly, the addition of 25 pM of SST at 1 mM glucose resulted in a 60% suppression of glucagon secretion in CTL mice (**Fig. 7b**). In HFD mice, glucagon secretion at 1 mM glucose was 100% higher than in CTL mice and the responses to exogenous somatostatin was markedly curtailed with no statistically significant inhibition of glucagon secretion detected (**Fig. 7b**). In isolated islets glucagon secretion in both CTL and HFD mice was suppressed by SST (25 pM), but was higher in HFD islets (**Fig. 7c**).

**Figure 7:**
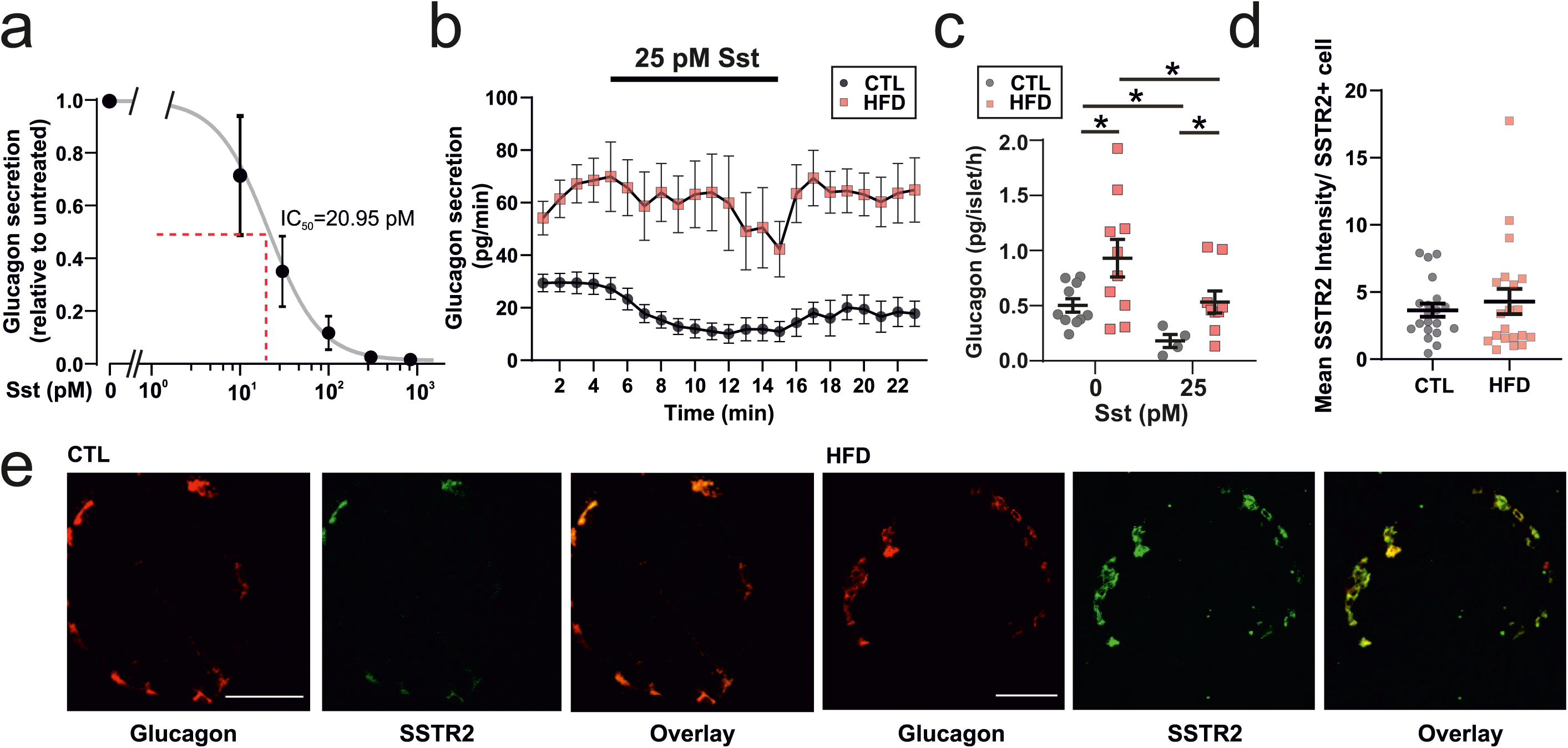
HFD results in changes in SST resistance. *a*. Dose response curve for SST on glucagon secretion measured in the perfusate from mouse pancreas in mice fed a chow diet. n=3 mice. Data is presented as mean±SEM. *b*. Glucagon measured in perfusate from CTL and HFD mice in response to 25 pM STT at 1mM glucose. n=9 mice in each group. Data is presented as mean±SEM. *c*. Glucagon secretion from isolated islets from 3 CTL and 3 HFD mice. T-test, p<0.05=*. Data is presented as mean±SEM. *d*. Staining of imbedded pancreata from CTL and HFD fed mice, for glucagon (red) and SSTR2 (green), scale bar is 50 µm. *e*. Analysis of SSTR2 staining intensity from *i*. Intensity was normalised to the number of cells expressing SSTR2. n=20 islets from 2 mice in each group. Data is presented as mean±SEM.

The above findings demonstrate that in response to HFD feeding, not only is SST secretion from delta-cells reduced, but alpha-cells also become resistant to SST. We reasoned that a reduction in SSTR2 expression may underlie the observed resistance. However SSTR2 staining in CTL and HFD pancreases revealed no differences (**Fig. 7d-e**), in line with recent reports from obese human donors [47].

## Discussion

Glucagon plays a significant role in the aetiology of diabetes [1] but little is known about how changes in alpha-cell function manifest in pre-diabetes. We have investigated alpha-cell function in response to prolonged ingestion of high amounts of fat (60% of total calories). We found that obese mice become hyperglucagonaemic and have impaired glucose-dependent inhibition of glucagon secretion. This echoes what is observed clinically in obese people [17, 18]. While it is hard to find commonalities in the literature regarding HFD-induced hyperglucagonaemia, our findings suggest that methodological aspects such as time of sampling have a strong influence on the level of circulating glucagon measured. Thus, these findings could potentially explain why measurements of plasma glucagon in HFD fed models differ markedly between studies [26-28, 48] and why it has been difficult relating them to human observations.

Glucagon secretion from alpha-cells depends on changes in [Ca^2+^]_i_. We now demonstrate that glucose regulates both the frequency and amplitude of [Ca^2+^]_i_ oscillations in alpha-cells. This is to our knowledge the first report distinguishing and analyzing these two components of the Ca^2+^ signal and demonstrating that glucose exerts statistically significant effects on these two key parameters in the regulation of glucagon secretion that have previously escaped detection. Unlike previous studies, this study has sufficient statistical power to detect an average change despite significant cell-to-cell variability; we have analyzed [Ca^2+^]_i_ in a much large number of alpha-cells (>500 for frequency and >120 for amplitude) than analyzed in earlier studies. Interestingly, it was not only the activity of single alpha-cells that changed with HFD feeding, the diet also changed the proportion of active alpha-cells. This could account at least for some of the change in frequency and indicates that the threshold for alpha-cell activity has been changed by HFD feeding. These findings suggest that several regulatory mechanisms underlie the changes in [Ca^2+^]_i_ and glucagon output observed with HFD feeding.

Alpha-cells are under strong paracrine regulation of SST [12, 14]. A recent study suggested that delta-cell [Ca^2+^]_i_ oscillatory activity is reduced after HFD feeding [49], in keeping with the reduction of somatostatin secretion we observe. We note that the effects of HFD on somatostatin secretion – when mice remain largely normoglycaemic – are different from those observed once hyperglycaemia has developed [50]. Not only was somatostatin secretion reduced in HFD mice, their alpha-cells were also much less sensitive to SST in the present study, with Ca^2+^ oscillation frequency persisting in HFD islet when exposed to SST. The cellular node of this resistance is not clear, although we can confidently say it is not due to a reduction in SSTR2 protein. In human alpha-cells, high concentrations of SST (30 nM) partially (70%) inhibits glucagon exocytosis [51]. Therefore, part of the resistance to SST in the HFD alpha-cells may be due to an effect on the sensitivity of the exocytotic machinery to SST. Intestinal D-cells secrete long-form SST (SST-28), which is distinct from that which is produced and secreted by pancreatic delta-cells (SST-14; see [45], [52] and [53]). As intestinal lipids stimulate GLP-1 and GIP release from the rat gut [54] and these hormones are elevated in rats fed a HFD [55], it is possible that gut SST-28 is similarly increased in response to high-fat feeding. An increase in circulating SST-28 may desensitise the alpha-cell SSTR2 receptor and/or exocytotic machinery to islet SST and result in the SST resistance we observe.

Although alpha-cells demonstrate SST resistance in response to HFD feeding, and SST secretion is reduced in 1 mM glucose, this cannot fully explain the increased alpha-cell [Ca^2+^]_i_ oscillation amplitude and glucagon secretion at 1 mM glucose, where there is very little SST secretion. (However, SST has recently been observed to tonically inhibit glucagon secretion at low (3.5 mM) glucose from the perfused rat pancreas [56]). As [Ca^2+^]_i_ amplitude was reduced in 1 mM glucose – something known to be under the control of intrinsic fuel-sensing mechanisms [11, 33] - we suggest that the hyperglucagonaemia present in 1 mM glucose may be driven by changes in alpha-cell metabolism. Fatty acid oxidation in alpha-cells has been shown to regulate the amplitude of [Ca^2+^]_i_ oscillations [33]. Therefore the hyperglucagonaemia observed at 1 mM glucose in HFD alpha-cells may be due to an increase in beta-oxidation, which is an important driver of glucagon secretion [24, 25].

The delta-cell is seemingly a key integrator and relay of circulating satiety and fuel signals to the islet alpha- and beta-cells, as it has been shown to express many GPCRs – including ghrelin receptors [57, 58] and leptin receptors [59]. The NEFA-responsive G-protein coupled receptor GPR120 is also expressed in delta-cells and its activation results in a reduction of SST secretion [60]. The elevation in circulating NEFAs in HFD fed mice would be expected to chronically reduce the output from delta- cells, explaining the hyperglucagonaemia observed under physiological glucose concentrations.

In conclusion, these findings link delta-cell dysfunction and SST resistance in alpha-cells directly to metabolic disease and demonstrated the importance of SST for the regulation of glucagon secretion in obesity and pre-diabetes.

## Figure Legends and Table

**Supplementary Figure 1:**
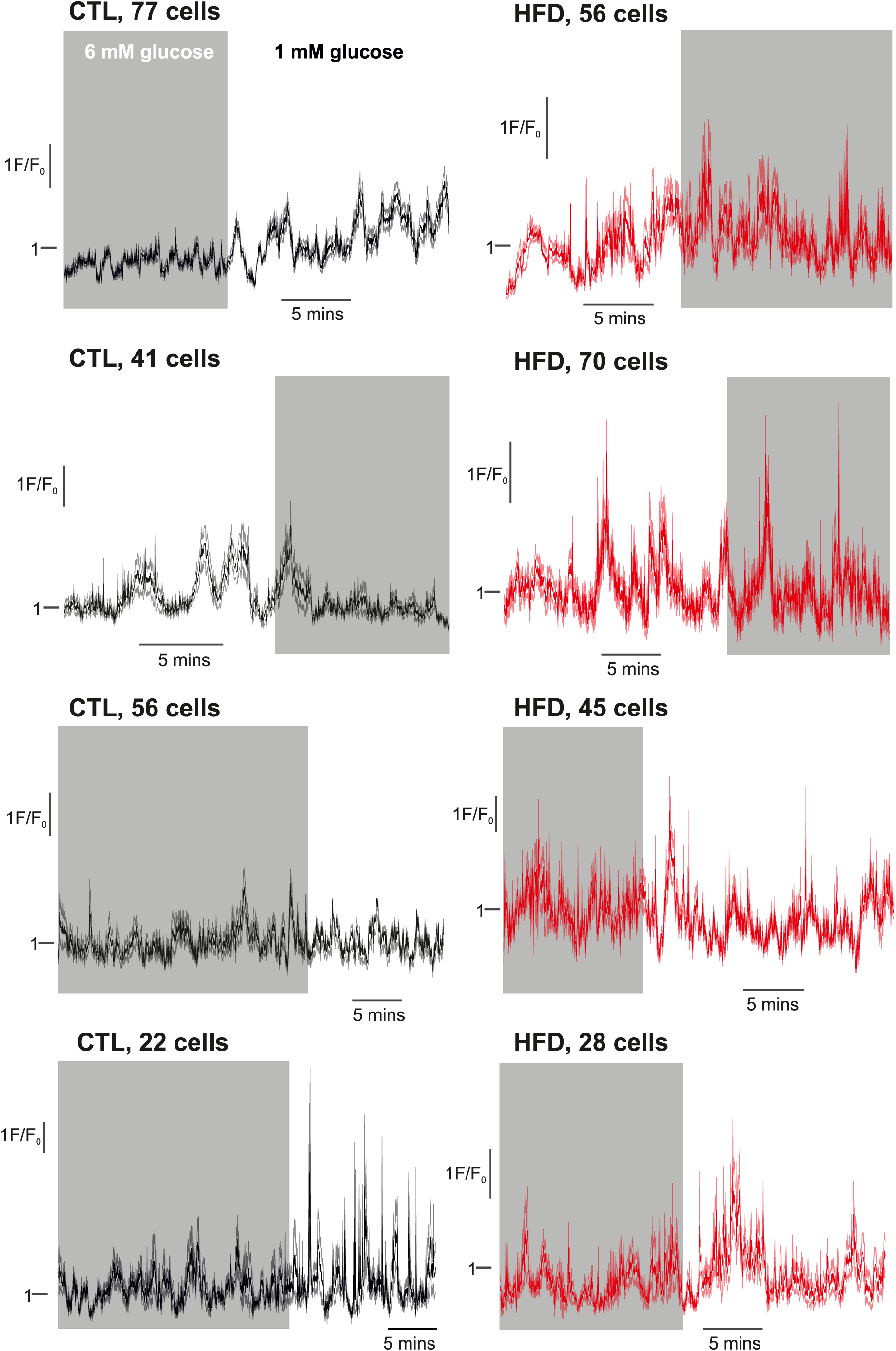
Whole-islet alpha-cell GCaMP3 oscillations in CTL and HFD mice. GCaMP3 was measured from islets from mice fed a CTL and HFD diet in response to 6 mM and 1 mM glucose (given in a randomized order). The GCaMP3 signal (F) from each individual alpha-cell was processed using `msbackadj` (MATLAB) and dividing by a period of baseline activity (F_0_). This was then averaged across all cells in the islet. Activity went through periods of being synchronised (on a ∼2 min timescale), as indicated by the waves in F/F_0_. Note that in response to 6 mM glucose, activity was suppressed in CTL islets but not in HFD islets. Data shown is mean±SEM from n=4 CTL mice and n=4 HFD mice.

## Acknowledgements

We thank Prof Leanne Hodson, Dr Siôn Parry and Prof David Hodson for feedback on this work, and Dr Claudia Guida with assistance in the RIA protocol. We also thank Prof Dame Fran Ashcroft for her help with EchoMRI measurements.

## Backmatter

### Competing interests

MvdB is an employee of Novo Nordisk. The other authors declare no competing interests.

### Data availability statement

All data is made freely available on reasonable request to the corresponding authors.

### Author contributions

LJBB, JGK and JAK designed the study and experiments. LJBB and JGK wrote the initial draft of the manuscript. JAK, TGH, NJGR, SA, LJBB and JGK conducted experiments and analysed data. All authors edited and approved the final version of the manuscript.

### Funding information

This study was funded by the following: Wellcome Senior Investigator Award (095531), Wellcome Strategic Award (884655), Sir Henry Wellcome Postdoctoral Fellowship (Wellcome, 201325/Z/16/Z) and a JRF from Trinity College. TH is supported by a Novo Nordisk postdoctoral fellowship run in partnership with the University of Oxford. JAK holds a DPhil from the OXION Programme (Wellcome). JGK held a fellowship with Novo Nordisk and now receives funding from Novo Nordisk Fonden.

